# aWCluster: A Novel integrative Network-based Clustering of Multiomics Breast Cancer Data

**DOI:** 10.1101/558700

**Authors:** Maryam Pouryahya, Jung Hun Oh, Pedram Javanmard, James C. Mathews, Zehor Belkhatir, Joseph O. Deasy, Allen R. Tannenbaum

**Affiliations:** Department of Medical Physics, Memorial Sloan Kettering Cancer Center, NY; Department of Medicine, Division of Endocrinology Diabetes and Bone Disease, Icahn School of Medicine at Mount Sinai, NY; Departments of Computer Science and Applied Mathematics & Statistics, Stony Brook University, NY

## Abstract

The remarkable growth of multi-platform genomic profiles has led to the multiomics data integration challenge. In this study, we present a novel network-based integration method of multiomics data as well as a clustering technique founded on the Wasserstein (Earth Mover’s) distance from the theory of optimal mass transport. We applied our proposed method of aggregating multiomics and Wasserstein distance clustering (aWCluster) to invasive breast carcinoma from The Cancer Genome Atlas (TCGA) project. The subtypes were characterized by the concordant effect of mRNA expression, DNA copy number alteration, and DNA methylation as well as the interaction network connectivity of the gene products. aWCluster successfully clusters the breast cancer TCGA data into classes with significantly different survival rates. A gene ontology enrichment analysis of significant genes in the low survival subgroup leads to the well-known phenomenon of tumor hypoxia and the transcription factor ETS1 whose expression is induced by hypoxia. In addition, immune subtype analysis in our clustering via aWCluster recovers the inflammatory immune subtype in a group demonstrating improved prognosis. Consequently, we believe aWCluster has the potential to discover novel subtypes and biomarkers by accentuating the genes that have concordant multiomics measurements in their interaction network, which are challenging to find without the network inference or with single omics analysis.

## Introduction

The molecular development of neoplasms occurs at a number of genomic, transcriptomic and epigenomic scales. The aggregation of multi-dimensional omics data potentially provides a comprehensive view of the etiology of oncogenesis and tumor progression at different molecular levels. Many sophisticated mathematical and statistical methods of multiomics data integration have been proposed, yet the need for an effective technique to improve the clinical outcome prediction remains a challenge in the era of precision medicine^1^. Large scale cancer genome projects, such as The Cancer Genome Atlas (TCGA) project include an unprecedented amount of multi-dimensional data to explore the entire spectrum of genomic abnormalities in human cancer^2^. With the exponential growth of such multi-dimensional data, the great need to obtain an integrated view of their interplay is becoming ever more pressing. A standard approach considers clustering each one-dimensional omics data separately, then finding the consensus of these clusterings^3^. Such methods result in loss of information and accumulation of noise in the database due to single omics treatment of data sets and not considering the impact of different layers on each other. A more elegant approach that we focused on in this work is integrating these multiomics in the very beginning followed by a single integrated clustering of the samples. Our method of aggregating multiomics and Wasserstein distance clustering (aWCluster) regards the gene expression of any transcript as a biological function of the gene copy number alteration and DNA methylation. Moreover, the gene expression can be modulated in any level by the other genes and their protein products in their interaction network. Our integrative measurements of genes in aWCluster, which is based on the interplay of these different layers in their interaction network, represent the functional concordance of these multiomics data.

Genomic instability can be caused by DNA copy number alteration and DNA methylation of CpG islands. In fact, amplification of oncogenes and deletion of tumor suppressors give rise to malignant neoplasms^4^. Furthermore, epigenetic aberrations caused by DNA methylation has a central role in tumor progression^5^. Combining the values of copy number alteration and DNA methylation with gene expression in a network-based manner significantly improves the statistical accuracy in characterizing the genes that have an essential role in cancer progression. Our method of integration in aWCluster is akin to CNAmet in aggregating the copy number, methylation, and gene expression data^6^. However, CNAmet does not consider the interaction network and does not include a clustering algorithm. We also include the continuous value of the multiomics data rather than the binary values {0,1} used in CNAmet to avoid the sensitive dependency on such thresholds.

In order to include the gene interaction network in aWCluster, we consider the weighted network for each sample as the result of an underlying stochastic process that is driven by interactions among connected nodes. For each sample, we construct a distribution of integrative omics measures across the nodes of the network, which is closely related to the invariant (stationary) measures of an associated Markov chain. We employ these integrative measures to cluster TCGA breast cancer data. Our clustering method in aWCluster is based on optimal mass transport (OMT) methods, utilizing the 1-Wasserstein distance^7^, also known as Earth Mover’s Distance (EMD), applied to the invariant distributions of integrative measures computed between samples^8^. Clustering via the Wasserstein distance in aWCluster is significantly more effective compared to clustering with Euclidean distance between samples. The invariant measure of the Markov chain considered in this work has an explicit closed formula which makes our method more efficient and interpretable than the spectral clustering methods used in other integrative methods such as similarity network fusion (SNF)^9^. Also, even though the iterative model of SNF is network-based, there was no consideration of the biological interaction network and the mechanistic interplay of multiomics layers. The difficulty of interpretability also occurs with the latent variables in matrix factorization methods such as iCluster^10^. One of the key advantages of aWCluster is its interpretability. Using the proposed aWCluster methodology, we are able to identify the significant genes in the specific cluster with a significantly low survival rate. We further perform gene ontology (GO) enrichment analysis on these genes utilizing a curated bioinformatics database (MetaCore) to discover the significantly correlated biological processes/pathways that could be related to the high mortality in this cluster.

The complexity of cancer etiology, the advent of large scale diverse genome-wide data, and the significant improvement in mathematical/statistical data analysis tools has resulted in considerable progress in the field of multiomics integration^11^. In addition, recent efforts in network-based analysis of ‘omics’ allow identification of new disease genes, pathways and rational drug targets that were not easily detectable by isolated gene analysis^12^. An effective bridge between the biology of cancer and mathematical techniques can bring about a comprehensive, meaningful and predictive model of multiomics integration with a medical target treatment plan for cancer patients. In this work, we strived to fulfill this goal and intertwine the mechanistic function of multiple regularity levels and the gene (protein) interaction network with advanced mathematical techniques for analyzing and clustering the data. The present study illustrates a novel mathematical way to integrate and cluster the multiple genomics data of TCGA breast cancer. Our clustering result is concordant with existing cancer subtypes, e.g., PAM50 in breast cancer, is effective in predicting survival rate, and facilitates the identification of the important driver genes in each sub-cluster, which in turn may allows researchers to glean new therapeutic approaches from the results.

## Materials and Methods

We adopted methods from OMT^7,13^ to measure the similarity of the integrative multiomics profiles between samples. To this end, we calculated the 1-Wasserstein distance (EMD) between the probability distributions of the integrative measures for the samples. More precisely, we first derived the integrative measures from the invariant measures of the stochastic matrix associated with the interaction network of genes. The integrative measures aggregate the gene expression, copy number alteration and methylation in a network-based fashion. We then utilized the Wasserstein distance to measure the similarity between every pair of probability measures *π*\*^1^ and *π*\*^2^ of integrative measures assigned to every two samples (Figure 1). Consequently, we applied these pair-wise distances (EMD) among samples to perform the hierarchical clustering of the data. In the following section, we first discuss our method of integration in aWCluster, then we review the clustering of samples via 1-Wasserstein distance on the gene interaction network.

**Figure 1.**
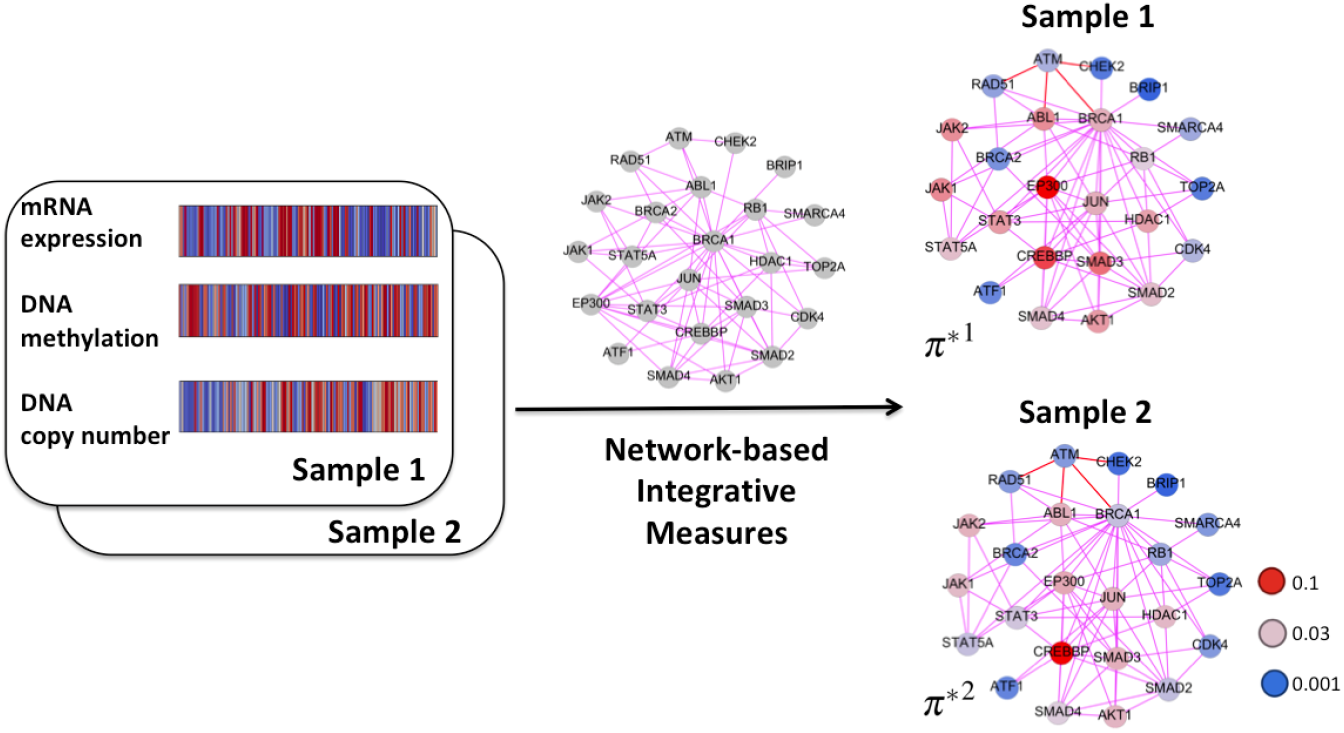
The integration of multiomics in aWCluster is network-based. We assigned an integrative measure to the nodes of the interaction network using the network connectivity and the values of gene expression, copy number alteration and methylation. These integrative measures are exploited by aWCluster to find the distances between samples and to cluster the sample space of TCGA breast cancer data. Here, we only show a small network for the purpose of illustration.

### Constructing Sample Specific Integrative Measures

We applied our method to integrate the multiomics data of TCGA breast carcinoma. The integrative measure is defined via the mRNA expression (z-score of RNA Seq V2 RSEM), copy number alteration (relative linear values from Affymetrix SNP6) and methylation (HM450) data of TCGA, and the interaction network from the Human Protein Reference Database (HPRD, http://www.hprd.org)^14^. After removing some missing data, our breast cancer TCGA data consists of mRNA expression of 18,022 genes for 1,100 cases, copy number values of 15,213 genes for 1,080 cases and methylation of 15,585 genes in 741 cases. The intersection of all three data resulted in 7,737 genes and 726 samples. Moreover, this gene set has 3,426 genes in common with HPRD in the largest connected component. Subsequently, we have all the three multiomics of 3,426 genes with their connected interaction network for 726 samples. We further considered making the method computationally less expensive by limiting the genes with the ones only in common with OncoKB (Precision Oncology Knowledge Base) genes (http://oncokb.org/) which is discussed extensively in the ‘Supplementary Information’ (SI) of the present paper.

We constructed a weighted graph by considering the given gene interaction network as a Markov chain^8^. Consider a gene *i* and its neighbor genes *j* ∈ *N*(*i*) in their interaction network (here in HPRD) for a given sample. Let *ge_k_* denote the expression level of gene *k* in a given sample. The principle of mass action allows us to compute the probability of the interaction of gene *i* to gene *j* (*p_ij_*) to be proportional to their expression, i.e. *p_ij_* ∝ (*ge_i_*)(*ge_j_*)^15^. By normalizing *p_ij_* such that such that *Σ_j_p_ij_* = 1, we have the stochastic matrix *p* of the Markov chain associated to the network as follows:

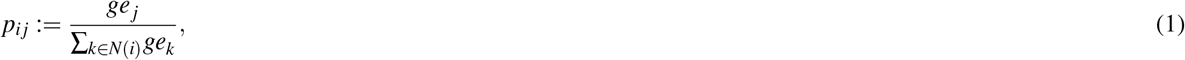

The Markov chain given by (1) reaches a stationary distribution which is invariant under a right multiplication by *p*, i.e. *πp* = *π*,^8^. Solving for the special stochastic matrix *p* defined by (1), *π* has the explicit expression:

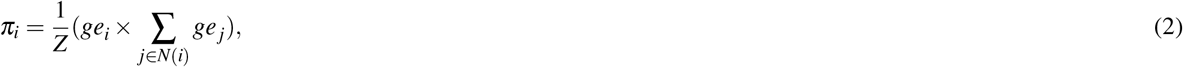

where *Z* is a normalization factor forcing *π* to be a probability vector. Note that this normalization is necessary since we want to consider the invariant measure to be a probability distribution over all genes for each specific sample. The invariant measure defined by (2), gives a value to each gene which is not only dependent on the gene expression of the gene *i*, but also on the total gene expressions of the neighboring genes *j* ∈ *N*(*i*).

We have extended this invariant measure to the integrative measure, which not only considers gene expression, but also copy number alteration and methylation. Our approach roughly follows that of CNAmet^6^, however, our method is network based. We characterize the genes that are upregulated with high expression, amplification and hypomethylation, and also have many connections with similarly upregulated genes in their interaction network. In fact, for TCGA data, the Spearman’s correlations between the gene expression and the copy number alteration across the samples are mostly significantly positive (see Figure S1 in the SI). Similar positive correlations exist between gene expression and 1–methylation (one *minus* methylation). Here, we considered the values of 1–methylation (which are positive as methylation values are between 0 and 1) since the high values of methylation have a reverse effect on the upregulation of genes. In CNAmet, the gene expression (mean across all samples) values have been multiplied by binary values assigned to copy number and methylation. In the proposed aWCluster method, we utilized the actual normalized values of copy number and methylation in the integrative measures for each sample.

More precisely, for the gene *i*, we define in an analogous manner to (2) the following:

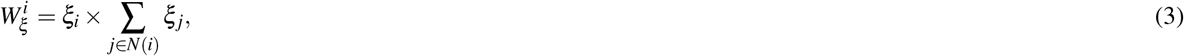

where *ξ* is any of the *ge, cn*, and *me*. In what follows, *W_ξ_* will denotes the corresponding the vector with components 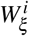. Similar to CNAmet, we introduce a scaling factor *ε*, which accounts for the concordance of copy number alteration and methylation. Yet again, we consider the actual normalized value of the data in the network-based fashion (Figure 1). For the gene *i*, the scaling factor is defined as follows:

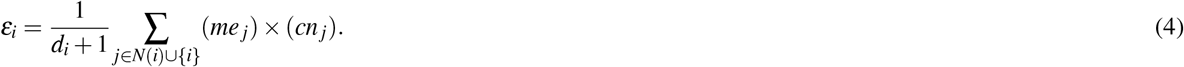

where *d_i_* is the node degree of the gene *i* in the interaction network. We next add the normalization factor *Z* in order to complete the formulation of the integrative measure as a probability vector. Consequently, we define the network integrative measure 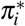 (for gene *i*) as follows:

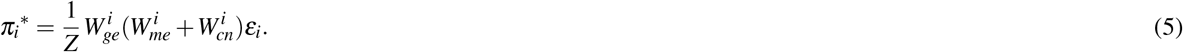

For each sample, we calculated a vector *π*\* = (*π_i_*\*)_*i*=1,…,*n*_ for all the *n* genes in our data sets. Here, we also present an alternative closed form formula for an integrative measure which is convenient for implementing. Let Adj denote the adjacency matrix of our network, i.e., the *n* × *n* matrix whose (*i*, *j*) entry is 1 if there is an edge (interaction) connecting node (gene) *i* and node *j*, and 0 otherwise. The vector *π*\* of integrative measures for a specific sample is defined as follows:

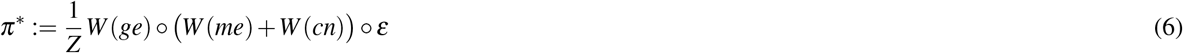

where

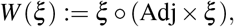

and

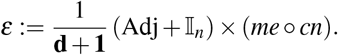

In these expressions, ○ denotes the component-wise (Hadamard) product, whereas × is the standard matrix multiplication. Also, **d** is the vector of all node degrees, **1** is all-ones vector, and 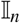 is the *n* × *n* identity matrix. Here, every row of the matrix 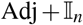 is divided by the corresponding component of **d +1**. Note that for each sample, we have defined a probability distribution *π*\* along the *n* genes taken as nodes of the corresponding network.

We finally applied methods from OMT^7^ to find the distance between a pair of vectors of the form *π*\* (in 6) assigned to every two samples. In fact, we measured the similarity between samples by finding the Wasserstein distance between the distributions of the integrative measures, which we define in more detail in the following section.

### Clustering via Wasserstein distance

Here, we use the theory of OMT to define distances among samples. OMT is a rapidly developing area of research that deals with the geometry of probability densities^7^. The subject began with the work of Gaspard Monge in 1781^16^ who formulated the problem of finding minimal transportation cost to move a pile of soil (“deblais”), with mass density *ρ*^0^, to an excavation (“remblais”), with a mass density *ρ*^1^. A relaxed version of the problem was introduced by Leonid Kantorovich in 1942^17^. Let *ρ*^0^, *ρ*^1^ ∈ *P*(Ω) where 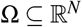 and *P*(Ω) = {*ρ*(*x*): ∫_Ω_*ρ*(*x*)*dx* = 1, *ρ*(*x*) ≥ 0}. The *1-Wasserstein distance*, also known as the *Earth Mover’s Distance (EMD*), is defined as follows:

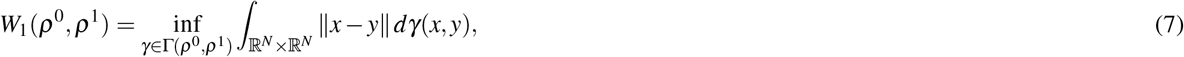

where Γ(*ρ*^0^, *ρ*^1^) denotes the set of all couplings between *ρ*^0^ and *ρ*^1^, that is the set of all joint probability measures *γ* on Ω × Ω whose marginals are *ρ*^0^ and *ρ*^1^ and the cost function of the transportation is defined as the ground distance *d*(*x*, *y*) = ∥*x* – *y*∥. We then used the pairwise EMD of samples for clustering our data. Each sample is represented as a vector of its pairwise distances to all other samples (its distance to itself is zero). We then applied the hierarchical agglomerative clustering to the sample vectors. Hierarchical clustering imposes a hierarchical structure on the samples and their stepwise clusters. To achieve a certain number of clusters, the hierarchy is cutoff at the relevant depth.

The aWCluster uses the theory of OMT to define distances among samples. OMT is a rapidly developing area of research that deals with the geometry of probability densities^7^. The subject began with the work of Gaspard Monge in 1781^16^ who formulated the problem of finding minimal transportation cost to move a pile of soil (“deblais”), with mass density *ρ*^0^, to an excavation (“remblais”), with a mass density *ρ*^1^. A relaxed version of the problem was introduced by Leonid Kantorovich in 1942^17^. Let *ρ*^0^, *ρ*^1^ ∈ *P*(Ω) where 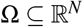 and *P*(Ω) = {*ρ*(*x*): ∫_Ω_*ρ*(*x*)*dx* = 1, *ρ*(*x*) ≥ 0}. The *1-Wasserstein distance*, also known as the *Earth Mover’s Distance (EMD*), is defined as follows:

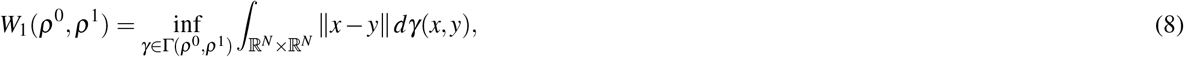

where Γ(*ρ*^0^, *ρ*^1^) denotes the set of all couplings between *ρ*^0^ and *ρ*^1^, that is the set of all joint probability measures *γ* on Ω × Ω whose marginals are *ρ*^0^ and *ρ*^1^ (see Figure S2 in the SI). Here, the cost function of the transportation is defined as the ground distance *d*(*x*, *y*) = ∥*x* – *y*∥.

An alternative and computationally more efficient formulation of the EMD (often called the Beckmann formulation^18^) is defined by optimizing the flux vector 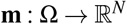 in the following manner:

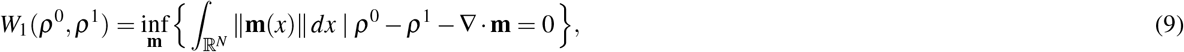

where ∥.∥ is the standard Euclidean distance based norm.

The optimization problem (8) has an analogous formulation on a weighted graph. Indeed, we consider a connected undirected graph 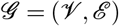 with *n* nodes in 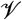 and *m* edges in 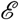. Given two probability densities 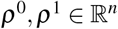 on the graph, the EMD problem seeks a joint distribution 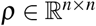 with marginals *ρ*^0^ and *ρ*^1^ minimizing the total cost Σ_*c_ij_ρ_ij_*_:

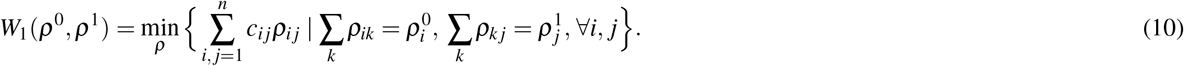

Here *c_ij_* is the cost of moving unit mass from node *i* to node *j* and is taken to be the minimum of the number of steps to go from *i* to *j*, namely, *c* is the ground metric on the graph. The minimum of this optimization problem defines a metric *W*_1_ (the Earth Mover’s Distance) on the space of probability densities on 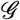. Note that this optimization problem consists of *n*^2^ variables.

Now we can give a graph-theoretic formulation of (9) as follows which gives an alternative way to compute the EMD on the graph 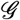:

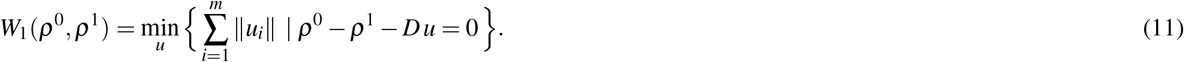

On the graph 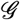, the *fluxes u_i_* are defined on the *m* edges, and 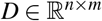 denotes the incidence matrix of 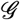 with an assigned orientation. More precisely, the incidence matrix *D* is a matrix with rows and columns indexed by the vertices and edges of *G* such that every entry (*i, k*) is equal to 1 if the vertex *i* is assigned to be the head of the edge *k* and is equal to −1 if it is the tail of *k*. Very importantly, note that the optimization problem in (11) depends on *m* variables, while the primal node based version of OMT on a graph in (10) depends on *n* × *n* variables. Thus formula (11) is certainly much more efficient especially when the graph is sparse, that is, *m* << *n*^2^.

In our computer implementations, we used the CVX optimization package^19^ to solve the optimization problem in (11). We used the pairwise EMD of samples for clustering TCGA data of breast cancer. Each sample is represented as a vector of its pairwise distances to all other samples (its distance to itself is zero). We then applied the hierarchical agglomerative clustering to the sample vectors. Hierarchical clustering imposes a hierarchical structure on the samples and their stepwise clusters. To achieve a certain number of clusters, the hierarchy is cutoff at the relevant depth. The optimal number of clusters can be determined with several techniques. In the present work, we validated the clustering via the following approach. First, we measured the homogeneity of clusters with silhouette mean values^20^. The optimal number of clusters is based on the silhouette values. The silhouette value for each sample is a measure of how similar that sample is to other samples in its own cluster compared to samples in other clusters. The *silhouette value* for the sample *i*, *s*(*i*) is defined as:

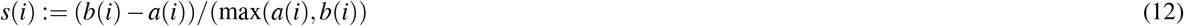

where *a*(*i*) denotes the average distance of the sample *i* to all samples within its own cluster (squared Euclidean distance between sample vectors), and *b*(*i*) denotes the minimum average distance of the sample *i* to samples of other clusters. Silhouette values range from −1 to +1. A high silhouette value indicates that the data are appropriately clustered. Therefore, we chose the optimal number of clusters by analyzing the average silhouette values of samples to make sure they stay close to 1.

For further evaluation of breast cancer clustering via aWCluster, we studied the difference of survival counts and survival rates among the clusters using the chi-squared test and log-rank test, respectively. We also compared our clustering results with the known PAM50 molecular subtypes of breast cancer (Luminal A, Luminal B, HER2-enriched, Basal-like, and Normal-like), which were first described in 2000 by Perou *et al*.^21–23^. This classification begins with the analysis of microarray expressions of 50 genes (known as the “PAM50” gene signatures) to cluster breast tumors into one of the subtypes.

In summary, aWCluster consists of three parts: integrating the multiomics, finding the Wasserstein distance among samples, and applying the hierarchical clustering to the samples space of the vectors of Wasserstein distances. In this work, we implemented aWCluster via MATLAB.

## Results

We primarily applied aWCluster to TCGA breast cancer data consisting of 3,426 genes and 726 samples. The visualization of the interaction network of these 3,426 genes is included in Figure S3 of SI. The hierarchical clustering of 726 samples of TCGA breast cancer via aWCluster is shown in Figure 2(a). The overall (ten-year) survival status in these clusters is given in the contingency Table 1. The p-value of the Pearson’s chi-squared test for this table is 0.02, indicating the significant separation of survival counts in our four breast cancer clusters. Moreover, the Kaplan-Meier plot of the clusters’ survival rate (truncated at 5 years) is illustrated in Figure 2(b). The log-rank test indicates a significantly different survival time between the four Kaplan-Meier curves (p-value=0.00187). We chose the number of clusters based on the silhouette values we discussed previously in the ‘Materials and Methods’ section. Here, choosing the three or four clusters had a very small effect on the silhouette values, however, clusters 1 and 2 (which are combined in the presence of three clusters) have very different survival rates. Therefore, we chose the number of clusters to be four. As shown in Figure 2(b), Cluster 2^*^ has a significantly low survival rate compared to the other three clusters. We note that excluding the samples with very short follow-up (less than 1 year) has minimal effect on the Kaplan-Meier curve (p-value < 0.01). The average days to last follow-up of the samples is 26 months.

**Table 1.** Overall (ten-year uncensored) survival status in the four clusters of 726 samples with breast cancer using 3,426 genes. The p-value of the chi-squared test for this contingency table is 0.02. The percentage of living and deceased cases in clusters are also included in parentheses.

| Survival | Cluster 1 | Cluster 2* | Cluster 3 | Cluster 4 | Total |
| --- | --- | --- | --- | --- | --- |
| Living | 39<br>(93%) | 42<br>(79%) | 319<br>(90%) | 231<br>(84%) | 628<br>(86%) |
| Deceased | 3<br>(7%) | 11<br>(21%) | 35<br>(10%) | 43<br>(16%) | 98<br>(14%) |
| Total | 42 | 53 | 356 | 275 | 723 |

**Figure 2.**
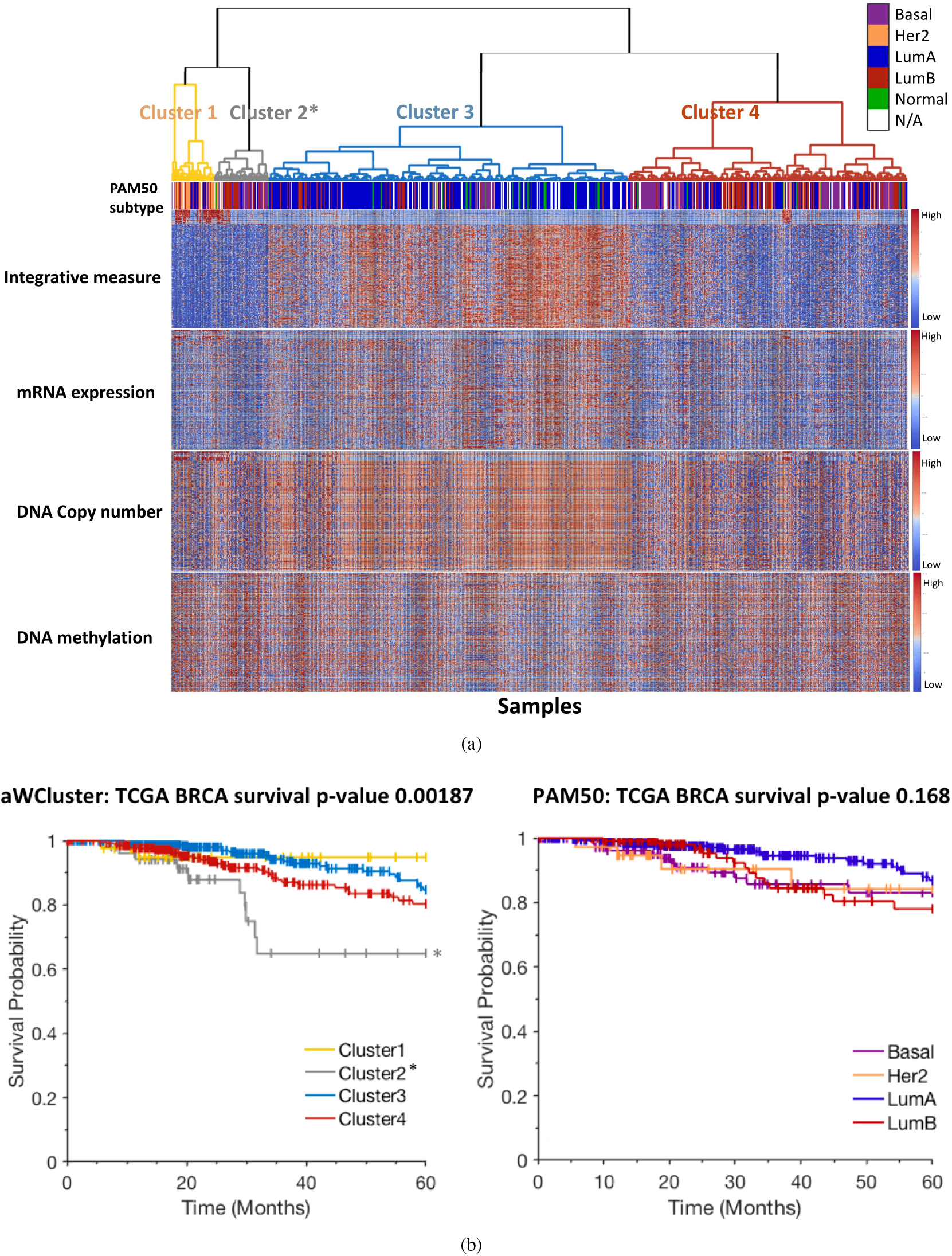
(a) Hierarchical clustering of 726 samples using 3,426 genes via aWCluster. The first bar underneath the hierarchical clustering corresponds to the PAM50 subtypes. The black color represents the samples whose PAM50 subtypes are not available. Furthermore, the heatmaps represent the integrative measures, gene expression, copy number alteration, and methylation of 150 genes selected based on ANOVA. Here, these top 150 genes are sorted (descending order) for cluster 1. The difference in the integrative measures between clusters is visually detectable in the heatmap. Of note, the samples are ordered based on clustering via integrative measures and if we started with one of these single omics we would not be able to achieve this clustering. We still see the pattern in the values of each single omics especially for gene expression and copy number alteration, however, they are not as clear as the one with the integrative values. (b) Graphical display of survival rate using the Kaplan-Meier curves with respect to PAM50 and aWCluster subgroups of TCGA breast carcinoma. Sample survival time (months) are plotted on the y-axis (truncated at 5 years), and the probability of survival calculated according to the Kaplan-Meier method is plotted on the x-axis. The p-value (log-rank test) for aWCluster is much lower than the PAM50 subtypes. The Normal subtype in PAM50 has been removed due to the small size (very minimal effect on p-value) and for better visualization. aWCluster also has a subgroup (Cluster 2*) with a significantly lower survival rate compared to the other three clusters.

We also compared our clustering with the well-known PAM50 molecular subtypes of breast cancer (Luminal A, Luminal B, HER2-enriched, Basal-like, and Normal-like). As shown in Table 2, our clustering substantially recovers the major PAM50 subtypes (chi-squared test’s p-value ≪ 10^−4^). We have the PAM50 subtype classification for 649 out of 726 samples in the database. Clusters 3 and 4 significantly distinguish Luminal A from the Basal-like subtype. Cluster 4 also includes most of the Luminal B subtype. Moreover, many of the Her2 enriched tumor types are in cluster 1 whereas the normal-like subtype is clustered together with Luminal A tumors in cluster 3. The set of 3,426 genes only includes 19 genes in common with PAM50 gene signatures. Also, as we expected, the overall survival status of cluster 3 which mainly consists of the Luminal A subtype is higher than cluster 4 which includes many of the Luminal B and Basal-like subtypes (p-value=0.014). Despite this consistency, our clustering results distiguished the subgroups with different survival rates much more significantly than the PAM50 subtypes (Figure 2(b)). Increasing the survival time in the Kaplan-Meier curve from 5 to 10 years improved the p-value for PAM50 subtypes to 0.06, albeit still not a significant value. However, the 10 year survival rate p-value (Kaplan-Meier curve) for aWCluster remains significant (p-value=0.004).

**Table 2.** aWCluster is significantly concordant with the PAM50 subtypes. The chi-squared test’s p-value ≪ 10^−4^ (even after excluding the normal subtype due to small counts). Cluster 3 includes most of the Luminal A subtype, whereas cluster 4 consists of the Basal-like and Luminal B subtypes.

| PAM50 | Cluster 1 | Cluster 2* | Cluster 3 | Cluster 4 | Total |
| --- | --- | --- | --- | --- | --- |
| Lum A | 7 | 8 | 240 | 83 | 338 |
| Lum B | 11 | 18 | 32 | 69 | 130 |
| Her 2 | 17 | 4 | 5 | 14 | 40 |
| Basal | 4 | 16 | 17 | 76 | 113 |
| Normal | 0 | 1 | 21 | 6 | 28 |
| Total | 39 | 47 | 315 | 248 | 649 |

We furthermore studied the immune subtypes of the TCGA breast cancer samples provided in the paper^24^. Table 3 provides the immune subtypes in our two largest clusters via aWCluster. As shown in Table 3, cluster 3 recovers most of the inflammatory immune subtype. This group was defined by Th17 and Th1 gene elevation and lower level aneuploidy^24^ and it has the best prognosis among TCGA data. Likewise, our cluster 3 which consists of mostly Luminal A, has a very good prognosis 2(b). Of note, cluster 3 includes even more samples of inflammatory immune subtype (81%) compared to the PAM50 Luminal A subtype (73%).

**Table 3.** Immune subtypes of breast cancer TCGA data in the two largest clusters (via aWCluster) provided by the paper^24^. Cluster 3 recovers most of the inflammatory immune subtype. This cluster shows a good prognosis. The percentages of each immune subtype in our clusters are also included in the table. The chi-squared test’s p-value ≃ 0 for this contingency table. The complete table including all four clusters is provided in ‘Additional data table S5’.

| Immune Subtype | Cluster 3 | Cluster 4 | Total (all four clusters) |
| --- | --- | --- | --- |
| Wound healing | 89<br>(40%) | 98<br>(44%) | 225<br>(31%) |
| IFN-γ dominant | 107<br>(39%) | 127<br>(47%) | 271<br>(36%) |
| Inflammatory | 113<br>(81%) | 21<br>(15%) | 139<br>(19%) |
| Lymphocyte depleted | 27<br>(44%) | 22<br>(36%) | 61<br>(8%) |
| TGF-β dominant | 18<br>(69%) | 5<br>(19%) | 26<br>(4%) |

Figure 2(a) shows the heatmap of integrative measures of 150 selected genes for 726 samples. The values of integrative measures are visually distinguishable between clusters which assures the accuracy of the clustering method. We utilized Analysis of Variance (ANOVA) to choose 150 top genes that have significantly different mean values of integrative measures across the four clusters. For better visualization, we further reordered these 150 genes based on the highest to lowest mean values in cluster 1. The highest values of integrative measures in cluster 1 (which has many HER2-enriched tumors) are by far ERBB2, GRB7, and PIK3C2A. These genes also have very significantly different mean values across the 4 clusters (after sorting based on ANOVA’s p-values). This result is consistent with the known co-amplification of ERBB2 and GRB7 in HER2-enriched breast cancer^25^. The complete list of these 150 genes has been provided in ‘Additional data table S1’ in the SI. We also provided the heatmap of gene expression, copy number alteration, and methylation in Figure 2(a). We still see the pattern in the values of each single omics. In fact, observing these patterns is expected, since the integrative measures are built of these single measures. However, these single omics patterns are not as clear as the one with the integrative values. It appears that the integrative procedure of aWCluster de-noises and smoothens the data and amplifies the contrast among extreme values of the genes in the single omics.

One of the advantages of aWCluster is its interpretability. The integrative measures of aWCluster are defined explicitly for all genes in the study, therefore, we can investigate genes that contribute the most in separating a specific cluster. Here, we are interested in the GO enrichment analysis of the significant genes in the clusters to see which biological processes/ pathways are related to the clusters with distinguished survival rates. As we see in Figure 2(b), cluster 2 has the lowest survival rate compared to all other clusters. This subgroup consists of 53 cases of mostly Luminal B and Basal-like subtypes. We identified the genes that have significantly different mean values in this cluster compared to the other three clusters using the t-test. We discovered that the significant genes genes in cluster 2 notably correspond with hypoxia. Tumor hypoxia, a well-known phenomenon where tumor cells have been deprived of oxygen, is a prominent issue in tumor physiology and cancer treatment^26^. Specifically, hypoxia appears to be strongly associated with tumor malignancy, resistance to treatment and the metastatic phenotype of cancer^27,28^. We chose 166 significant genes in cluster 2 based on the (t-test) Bonferroni corrected p-value less than 0.01 (Additional data table S2). The GO enrichment analysis of these genes was performed via MetaCore software (Thomson Reuters). MetaCore is an integrated software based on a manually-curated database of molecular interactions, molecular pathways, gene-disease associations, chemical metabolism, and toxicity information. The top ten biological processes presented with very small p-values (≪ 10^−6^) are shown in Table 4. These p-values correspond to the hypergeometric test performed by MetaCore using the number of input genes, ontology related genes, and the total number of genes in the database. Here, the top biological processes (first and second and also fifth) strongly correspond to hypoxia (p-value ≪ 10^−6^) which is a central issue in tumor physiology and cancer treatment. The protein/gene interaction network of these 166 genes is also presented in Figure 3. As we see in this figure, the network is very much connected which indicates that many of these 166 genes are related to each other. The three hub nodes, ETS1, AP-1, and STAT3 located within the nucleus are highly connected to many other proteins in the network. We will discuss these proteins and the genes related to them in more detail in the discussion section. We similarly chose the significant genes in the other three clusters using the t-test’s p-value, however, hypoxia did not appear as a significant top biological process in the other GO enrichment analysis.

**Table 4.** Top 10 biological processes obtained from the gene ontology enrichment analysis of the significant genes for the subgroup with the worst survival outcome (via MetaCore). Top biological processes (first, second and also fifth) are strongly correlated with hypoxia.

|  | Biological Processes |
| --- | --- |
| 1 | Response to decreased oxygen levels |
| 2 | Response to hypoxia |
| 3 | Regulation of cellular biosynthetic process |
| 4 | Cellular macromolecule metabolic process |
| 5 | Response to oxygen levels |
| 6 | Regulation of macromolecule biosynthetic process |
| 7 | Regulation of biosynthetic process |
| 8 | Positive regulation of macromolecule biosynthetic process |
| 9 | Positive regulation of cellular biosynthetic process |
| 10 | Positive regulation of nucleobase-containing compound metabolic process |

**Figure 3.**
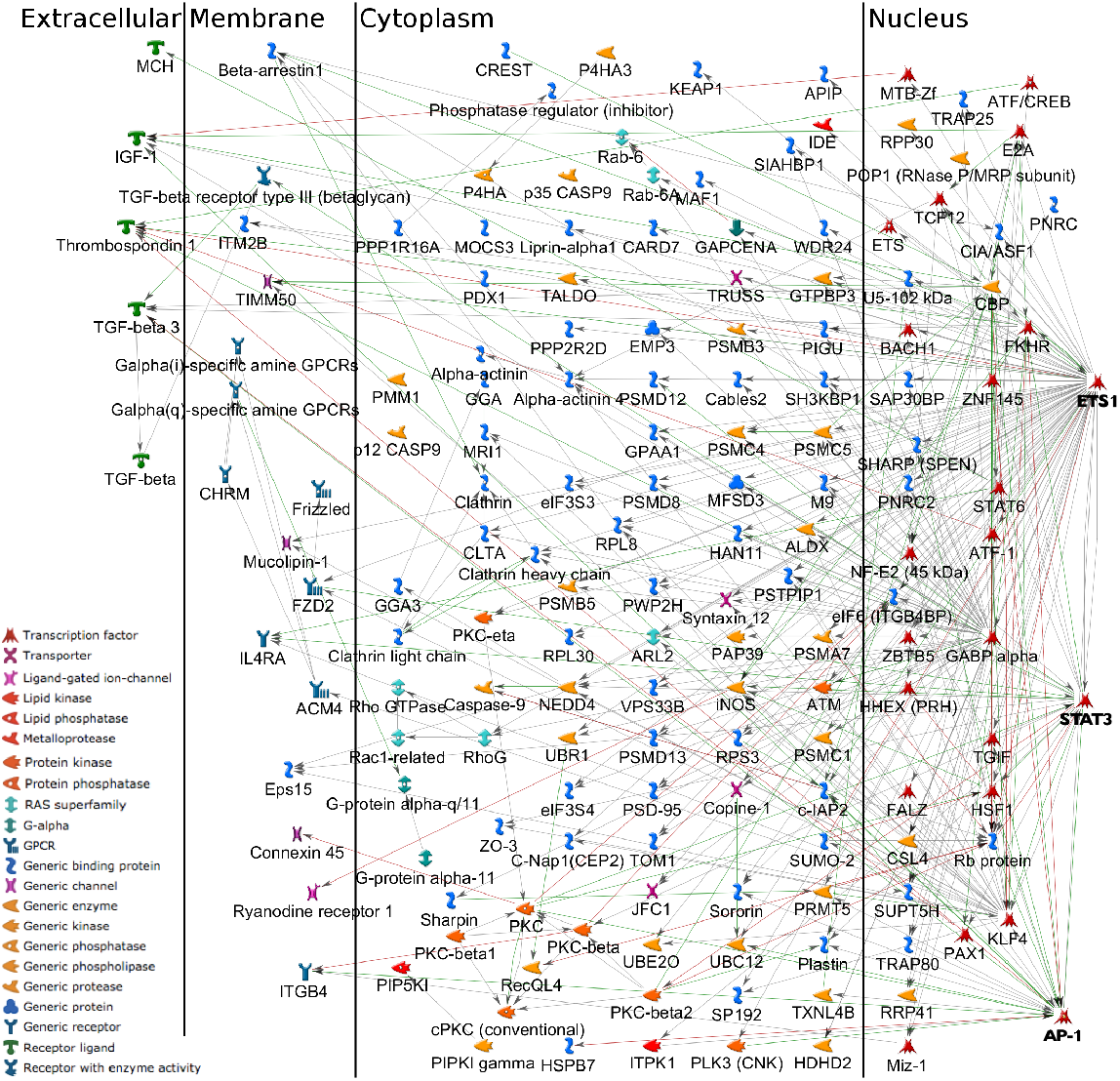
The protein (gene) interaction network of the significant genes for the subgroup with the worst survival outcome (via MetaCore). The network includes the hub nodes ETS1 as well as AP-1 and STAT3 located within the nucleus.

We finally used aWCluster to cluster TCGA breast cancer data using the genes that are also included in the OncoKB database. This smaller network with 290 genes is computationally much more efficient. The clustering of the 726 samples using this gene set is significantly consistent with our previous clustering using the 3,426 genes (chi-squared test’s p-value ≪ 10^−4^ for contingency Table S3) indicating the robustness of aWCluster’s methodology. We included the details of this clustering along with more clinical outcome analysis in the SI.

In order to investigate the importance of integrating the multiomics in aWCluster, we repeated all the steps of aWCluster with only the gene expression data (similarly with only copy number alteration/ methylation). The invariant measures given in (2) only depend on the gene expressions. After excluding copy number alteration and methylation, the Kaplan-Meier curves of clusters do not have a significant survival rate difference where the log-rank’s p-value is 0.09. Even though the difference in survival counts between two major clusters (4 and 5) is still significant (chi-squared test’s p-value= 0.02), it is not as strongly separated as in the integrative method (p-value= 0.014). Considering the copy number alteration or methylation solely results in even worse clustering in terms of survival difference between clusters. Therefore, the integrative procedure of aWCluster is crucial in the clustering of TCGA data.

## Discussion

The effective integration of large-scale cancer genome data can significantly improve clinical outcome predictions. Aggregating multiomics data leads to detection of biomarkers that are perturbed in a certain subpopulation of cancer patients. In addition, cancer-related genes do not act individually, but within an interaction network, which must be explicitly incorporated into the integration procedure. Therefore, network based integration of all the genomic, transcriptomic, and epigenomic data increases the information content and accuracy of the results more than any of the single omics studies separately. We propose the use of aWCluster for integrating the multiomics and clustering samples (patients). The network based integration of aWCluster considers the mRNA expression, DNA methylation, and copy number alteration of the genes in their corresponding neighborhoods of the interaction network. The concordance of the mutiomic values for a gene and connectivity to many similar genes results in a high integrative measure of the gene. aWCluster defines the similarity measured among samples by the Wasserstein distance between the distributions of the samples of the integrative measures along the genes. Applying aWCluster to TCGA breast cancer data successfully clusters data with significantly different survival rates. Moreover, we were able to recover the well-known PAM50 subtypes. aWCluster of breast cancer TCGA data also identified the inflammatory immune subtype which has an improved prognosis. The immune subtype analysis of TCGA data has been done through the characteristic immuno-oncologic gene signatures^24^. The identification of the inflammatory immune subgroup with increased survival in our independent clustering results may support the existence of other gene signatures in this subtype. Our clustering results also reveal a subgroup of breast cancer patients with substantially poor survival outcome. We performed the GO enrichment analysis on the genes that have significantly different values in this cluster compared to others. The analysis discovered that this gene set is significantly related to the biological process of hypoxia. Also, as we see in Fig. 3, the network of these genes is very densely connected, with a hub node ETS1 which is a transcription factor included in the list of 166 significant genes of the subgroup with the worst survival rate.

Tumor hypoxia, a well-known phenomenon where tumor cells have been deprived of oxygen, is a prominent issue in tumor physiology and cancer treatment^27^. Notably, tumor hypoxia is strongly associated with tumor propagation, malignant progression, and tumor angiogenesis, with cells in hypoxic regions of a tumor more resistant to the effects of radiation and chemotherapy^29^. In hypoxic environments, cellular adaptive processes are provoked which are mediated by the elevated expression of hypoxia-inducible genes thereby enabling tumor cells to adapt and survive under harsh conditions^30^. The hypoxia-inducible factor (HIF-1) is the principal transcription factor related to hypoxia, and it has been demonstrated that HIF-1 activity is increased in various tumors relative to that found within normal tissues^31,32^. Along with HIF-1, members of the v-ets erythroblastosis virus E26 oncogene homolog (ETS) family transcription factors, most prominently the proto-oncogene ETS1, participate in the upregulation of hypoxia-inducible genes^30^. Oikawa *et al*.^26^ showed for the first time that ETS1 is induced in the setting of hypoxia via the transcriptional activity of HIF-1. Of note, increased expression of ETS1 is seen in a variety of solid cancers including lung, colorectal, sarcoma, and squamous cell carcinomas, and higher levels have correlated with a higher incidence of lymph node metastasis and overall worse prognosis^33^. This gene is also involved in tumor progression in breast cancer, where in the setting of hypoxia, increased expression by mammary epithelial cells contributes to aggressive tumor phenotypes by activating the transcription of genes involved in angiogenesis, extracellular matrix remodeling, cell adhesion, and invasion^26,34^. Moreover, increased expression of ETS1 is associated with increased risk of recurrence and worse prognosis in human breast cancers^35^.

In addition, Activator Protein 1 (AP-1), another hub node in our network of significant genes (Figure 3) has also been identified as a hypoxia-inducible transcription factor^36^. C-jun, a proto-oncogene, encodes a major component of AP-1 transcription factors, which are key regulators of immediate-early signals directing cellular proliferation, differentiation, survival, and environmental stress responses^37^. AP-1 appears to be involved in the modulation of the apoptotic pathway and also plays a protective role in cellular response to DNA damage^38,39^. Piret *et al*.^40^ demonstrated an AP-1 mediated protective role of hypoxia against cell death induced by the chemotherapeutic agent etoposide. Similarly, in the setting of hypoxia, an anti-apoptotic role of AP-1 was seen in paclitaxel exposed breast cancer cells^41^.

Furthermore, STAT3 is another hub node in our network and is included in the set of 166 significant genes related to the subgroup with the worst survival. Hypoxia can induce the activation of transcription 3 (STAT3) protein, with the hypoxia-induced biochemical alterations likely contributing to drug resistance under hypoxic conditions^42^. Notably, hypoxia-induced STAT3 accelerates the accumulation of HIF-1 protein and has been shown to prolong its half-life in solid tumor cells^43^. Moreover, in a triple-negative breast cancer cell line, STAT3 has been shown to play a key role in hypoxia-induced chemoresistance to the chemotherapeutic agent cisplatin^44^.

Employing aWCluster, we combined the mRNA expression, DNA methylation and DNA copy number of alteration in a network-based fashion. We used the theory of optimal mass transport to find the clusters of samples with concordant genes’ integrative measures. One important advantage of our approach is preserving the integrative measure for all the genes, thereby enabling us to identify significant genes in each cluster.

We should note that our methodology does not identify the dominant etiology of the hypoxia subtype. In particular, these cases may represent an evolution from states represented by another cluster. Another limitation of the present work is that we only consider an undirected form of the interaction network. In addition to considering directionality, we previously studied the type of control (e.g., activator and repressor in transcription networks) in biological networks^45^. Accordingly, in future work, we plan to include the effect of both the network direction and regulation control type in our integrative measures. We would also like to consider more general versions of OMT^46^ to directly work with the vector-valued data.

Finally, we are also interested to integrate omics data with other data types such as imaging and drug response data^47^. In fact, the general framework of the methodology proposed in this work can be applied to other multi-dimensional data. After an appropriate integration of the data such as we did in this work, we can apply aWCluster for clustering and also discovering important features in the data of interest. We believe that the integration of multiomics/ biological data paves the way for precision medicine in treating sophisticated diseases such as cancer. To this end, we proposed a novel integrating method, based on fundamental optimal mass transport considerations, that allows for the interactive relationship among different omics layers, and accurately clusters breast cancer samples with significantly different survival rates.

## Acknowledgements

This study was supported by AFOSR grant (FA9550-17-1-0435), a grant from National Institutes of Health (R01-AG048769), MSK Cancer Center Support Grant/Core Grant (P30 CA008748), and a grant from Breast Cancer Research Foundation (BCRF-17-193).

## Additional Information

**Supplementary Information** accompanies this paper.

