## Supplementary Information for "aWCluster: A Novel integrative Network-based Clustering of Multiomics Breast Cancer Data"

We applied the aWCluster method to cluster TCGA breast cancer data of 726 samples. To this end, we primarily focused on multiomics of the large set of 3,426 genes which is discussed in the main paper. Here, we made the method computationally more efficient by limiting the genes to only those in common with OncoKB. OncoKB (http://oncokb.org/) currently has 1020 cancer-associated genes including the MSK-IMPACT genes (https://www.mskcc.org/msk-impact). Our gene set of 3,426 genes has 290 genes in common with OncoKB in the largest connected component of the HPRD network. As noted in the **Results** section, the effect of this dimension reduction of genes on the clustering is not statistically significant and therefore is more computationally desirable.

The interaction network is derived from HPRD by finding the largest connected component of the genes in the intersection of the HPRD and TCGA databases. The network visualization of these 290 genes is included in Figure S4. The clustering of the 726 samples using this gene set is significantly consistent with our previous clustering using the 3,426 genes (chi-squared test's p-value $\simeq 0$ for contingency Table S1 ) indicating the robustness of aWCluster's methodology. Similar to the clustering via 3,426 genes, the hierarchical clustering of 726 samples of TCGA breast cancer using these 290 genes results in two major clusters of 348 cases and 227 ones. The two clusters have a significantly different survival count as shown in the contingency table included in Figure S5(a). The p-value for the chi-squared test of this table is 0.001. The log-rank test indicates a significantly different survival time between the two Kaplan-Meier curves with p-value=0.001 (Figure S5(b)). Similar to clustering via 3,426 genes, our clustering using 290 genes is significantly concordant with the PAM50 subtypes (Table S2), despite having only 5 genes in common with PAM50 gene signatures. The OncoKB gene set consists of known cancer-associated genes which are linked to key cancer-related biological processes/ pathways. Therefore, using this gene set may not be appropriate to provide novel information via the GO enrichment analysis.

We further explored the clinical data of the TCGA breast cancer samples in these clusters. We investigated the tumor stages, tumor sizes and metastases of the samples (Table S3 and S4). Most of the tumors are in stage II (410 of 720 samples). We restricted the samples to only stage II tumors to investigate whether we still have a significant survival rate difference between cluster 1 and cluster 2. As shown in the Kaplan-Meier curves of Figure S6, the survival time is still significantly higher in cluster 1 (log-rank test's p-value= 0.01). Also, the TNM analysis of the tumors in the two clusters reveals that proportionally the tumors in cluster 2 are larger than the tumors in cluster 1 (chi-squared test's p-value= 0.02 for the tumor size contingency table (for cluster 1 and 2) in S4. This is consistent with the fact that cluster 1 (which has smaller tumors) has a better survival rate.

**Table 1.** We compared the sample cluster membership via aWCluster using 290 genes and 3,426 genes. Two major clusters are significantly consistent with the chi-squared test's p-value of $\simeq 0$ for their contingency table.

| aWCluster | Cluster 3 (via 3,426 genes) | Cluster 4 (via 3,426 genes) | Total |
|---|---|---|---|
| Cluster 1 (via 290 genes) | 304 | 44 | 348 |
| Cluster 2 (via 290 genes) | 51 | 164 | 215 |
| Total | 355 | 208 | 563 |

**Table 2.** aWCluster using 290 genes (in common with OncoKB) substantially recovers the major PAM50 subtypes. The chi-squared test's p-value $\simeq 0$ for this contingency table. Clusters 1 and 2 significantly distinguish Luminal A and Basal-like subtypes (p-value $\simeq 0$). Many Her2 subtypes are in Clusters 3 and 4.

| PAM50 | Cluster 1 | Cluster 2 | Cluster 3, 4 | Cluster 5 | Total |
|---|---|---|---|---|---|
| Lum A | 233 | 68 | 8 | 29 | 338 |
| Lum B | 35 | 57 | 9 | 29 | 130 |
| Her 2 | 2 | 14 | 18 | 6 | 40 |
| Basal | 15 | 63 | 5 | 30 | 113 |
| Normal | 21 | 3 | 1 | 3 | 28 |
| Total | 306 | 205 | 41 | 97 | 649 |

**Table 3.** Tumor stages of clusters (using the 290 genes in common with OncoKB) are presented for 720 samples (due to the missing values for some samples). The majority of the tumor samples (410 samples) are stage II.

| Tumor Stage | Cluster 1 | Cluster 2 | Cluster 3, 4 | Cluster 5 | Total |
|---|---|---|---|---|---|
| Stage I | 66 | 29 | 6 | 10 | 111 |
| Stage II | 189 | 132 | 26 | 63 | 410 |
| Stage III | 87 | 58 | 14 | 28 | 187 |
| Stage IV | 3 | 4 | 0 | 1 | 8 |
| Stage X | 1 | 2 | 0 | 1 | 4 |
| Total | 346 | 225 | 46 | 103 | 720 |

**Table 4.** Tumor sizes and metastases of samples in each cluster (using the 290 genes in common with OncoKB). Tumor sizes are presented for 724 samples and metastases are included for 589 samples (due to the missing values for some samples). Proportionally the tumors in cluster 2 are larger than the tumors in cluster 1 (chi-squared test's p-value= 0.02 for the tumor size contingency table including only clusters 1 and 2).

| Tumor Size | Cluster 1 | Cluster 2 | Cluster 3, 4 | Cluster 5 | Total |
|---|---|---|---|---|---|
| $T_1$ | 103 | 48 | 14 | 17 | 182 |
| $T_2$ | 183 | 138 | 26 | 71 | 418 |
| $T_3$ | 57 | 28 | 5 | 12 | 102 |
| $T_4$ | 5 | 12 | 1 | 4 | 22 |
| Total | 348 | 226 | 46 | 104 | 724 |

| Metastases | Cluster 1 | Cluster 2 | Cluster 3, 4 | Cluster 5 | Total |
|---|---|---|---|---|---|
| $M_0$ | 274 | 189 | 35 | 81 | 579 |
| $M_1$ | 3 | 5 | 0 | 2 | 10 |
| Total | 277 | 194 | 35 | 83 | 589 |

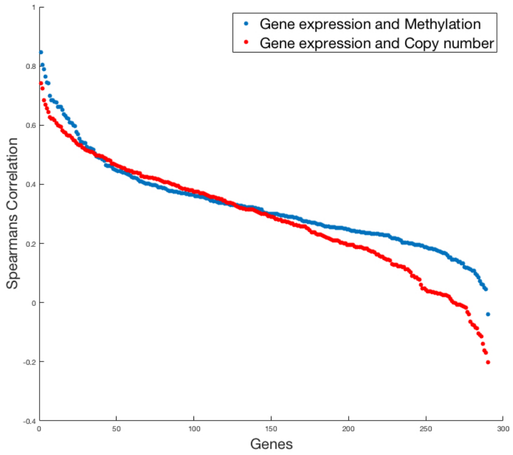


**Figure 1.** Spearman's correlation between gene expression and copy number alterations/ methylation for TCGA breast cancer data. Here, we considered 290 genes in common between TCGA, OncoKB and HPRD for 726 samples. The correlations between gene expression and copy number alteration are mostly positive (red plot). Similarly the correlations between gene expression and 1−methylation are positive (blue plot). Therefore, we considered the values of 1-methylation in our integrative formula of aWCluster.

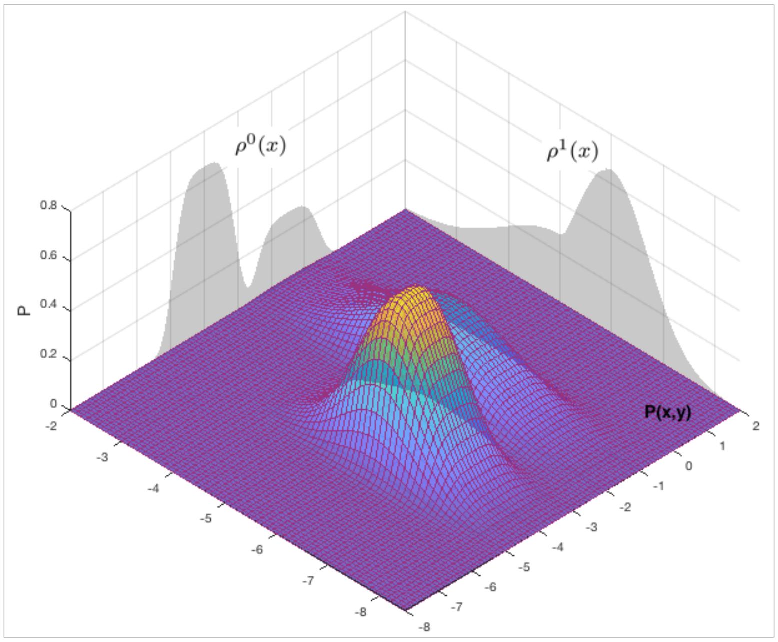


**Figure 2.** Kantorovich proposed using the couplings between $\rho^0$ and $\rho^1$ in the original Monge problem. This allows one to employ linear programming to solve the problem.

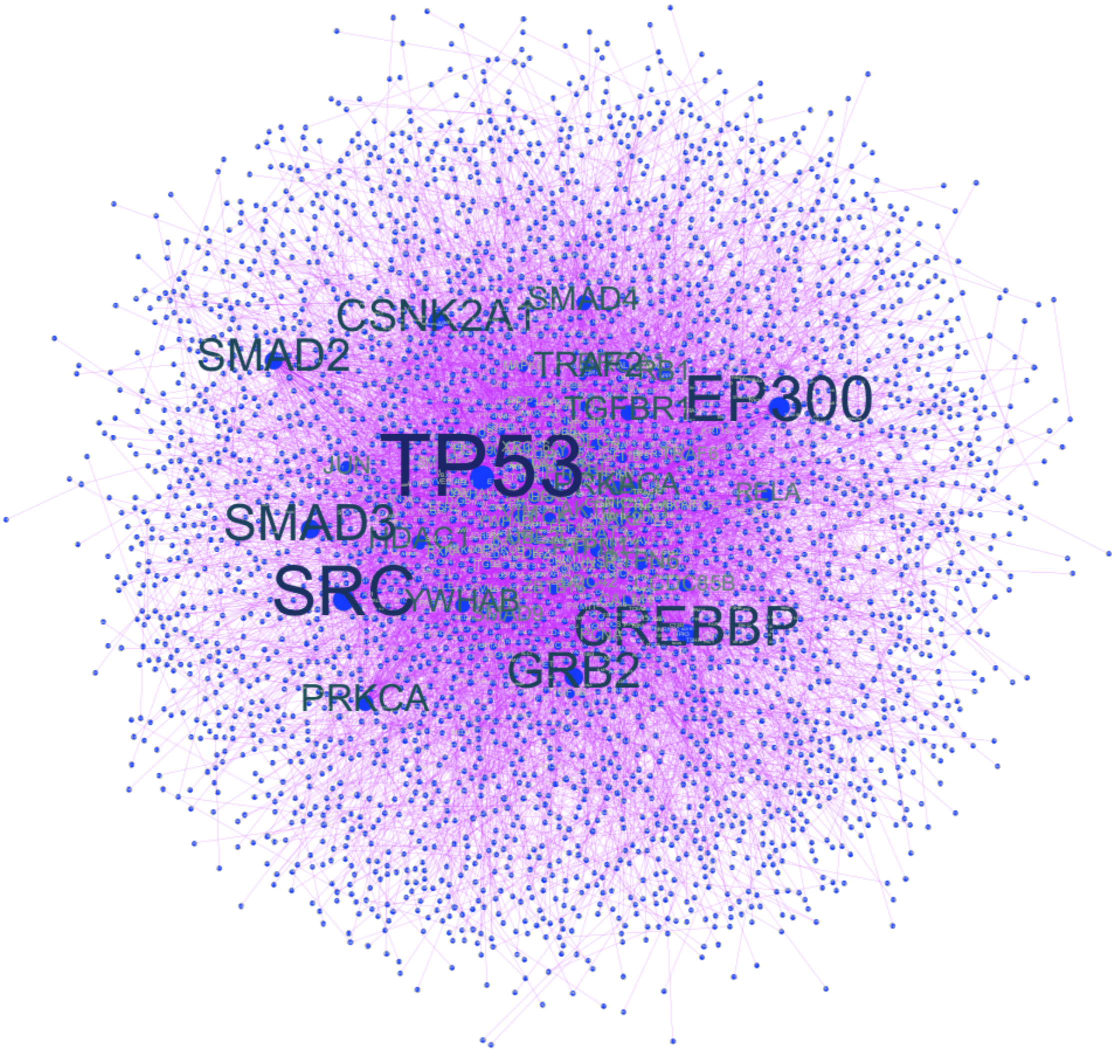


**Figure 3.** The network visualization (via Gephi). The network consists of 3,426 nodes (genes) and 9,385 edges. The sizes of the nodes (and font's size of node labels) are proportional to the node-degree. In aWCluster, we defined an integrative measures for the nodes of this network for each sample. We then used this weighted network for clustering of the sample space.

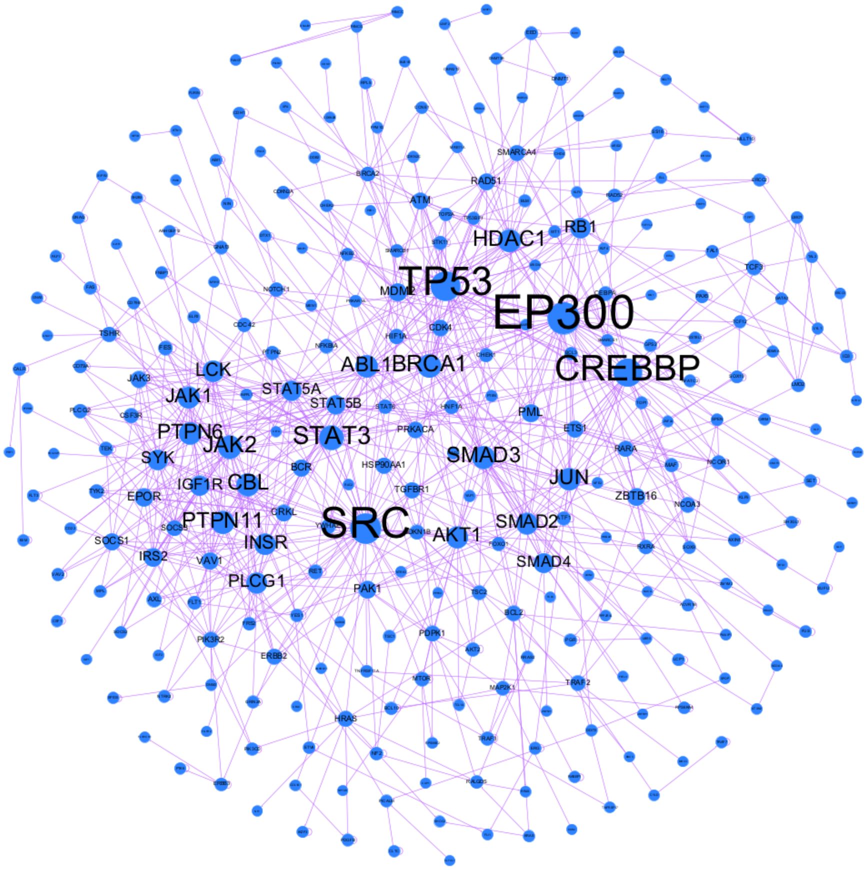


**Figure 4.** The network visualization (via Gephi). The network consists of 290 nodes (genes) and 824 edges. The sizes of the nodes (and font's size of node labels) are proportional to the node-degree. In aWCluster, we defined an integrative measures for the nodes of this network for each sample. We then used this weighted network for clustering of the sample space.

**Table 5.** Immune subtypes of breast cancer TCGA data in four clusters (via aWCluster). Cluster 3 recovers most of the inflammatory immune subtype. This cluster shows a good prognosis. The percentages of each immune subtype in our clusters are also provided in the table.

| Immune Subtype | Cluster 1 | Cluster 2* | Cluster 3 | Cluster 4 | Total |
|---|---|---|---|---|---|
| Wound healing | 19 | 19 | 89 | 98 | 225 |
| | (8%) | (8%) | 40%) | (44%) | (31%) |
| IFN-$\gamma$ dominant | 15 | 22 | 107 | 127 | 271 |
| | (5%) | (8%) | (39%) | (47%) | (36%) |
| Inflammatory | 1 | 4 | 113 | 21 | 139 |
| | (0.7%) | (3%) | (81%) | (15%) | (19%) |
| Lymphocyte depleted | 6 | 6 | 27 | 22 | 61 |
| | (1%) | (1%) | (44%) | (36%) | (8%) |
| TGF-$\beta$ dominant | 1 | 2 | 18 | 5 | 26 |
| | (4%) | (8%) | (69%) | (19%) | (4%) |
| Total | 42 | 53 | 354 | 273 | 722 |
| | (6%) | (7 %) | (49 %) | (38 %) | |

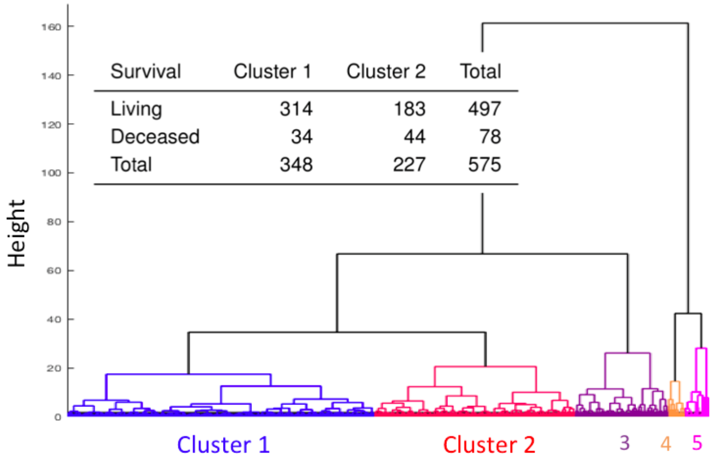


(a)

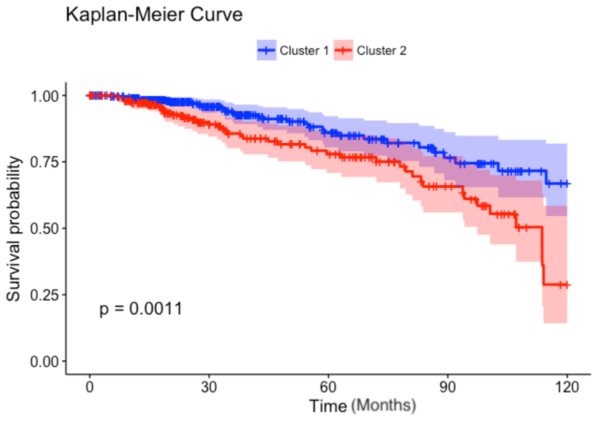


(b)

**Figure 5.** (a) Hierarchical clustering of 726 samples using 290 genes via aWCluster. The contingency table of survival counts shows a significant separation in two major clusters. (b) Graphical display of survival rate using the Kaplan-Meier curves. Sample survival times (months) are plotted on the x-axis (truncated at 10 years), and the probability of survival calculated according to the Kaplan-Meier method is plotted on the y-axis. The shaded areas illustrate the 95% confidence intervals.

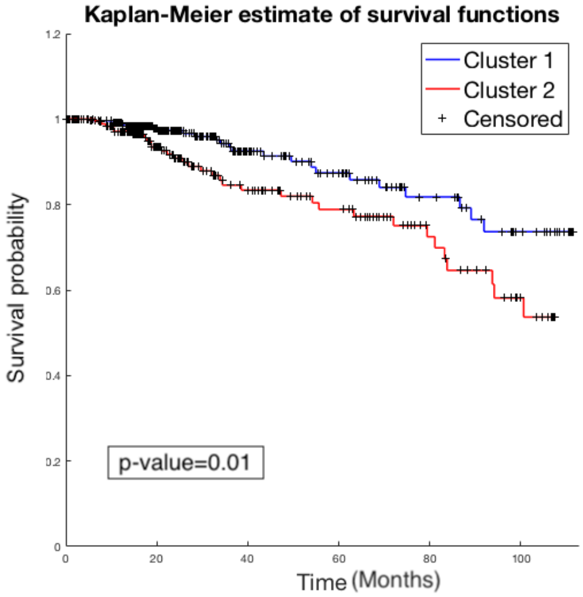


**Figure 6.** After restricting the samples to only stage II tumors, we still have significant survival rate difference between cluster 1 and cluster 2. As shown in the Kaplan-Meier curves, the survival rate is still significantly higher in cluster 1 with p-value= 0.01 for the log-rank test.

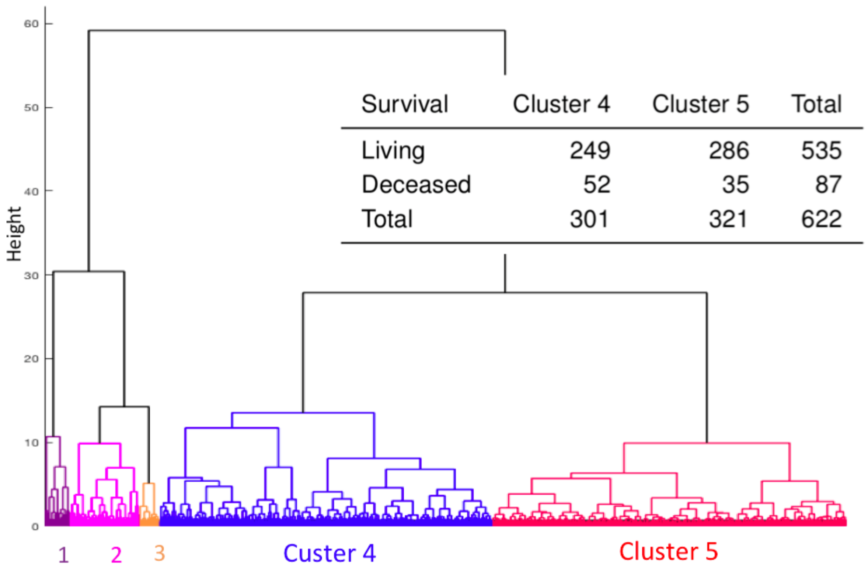


(a)

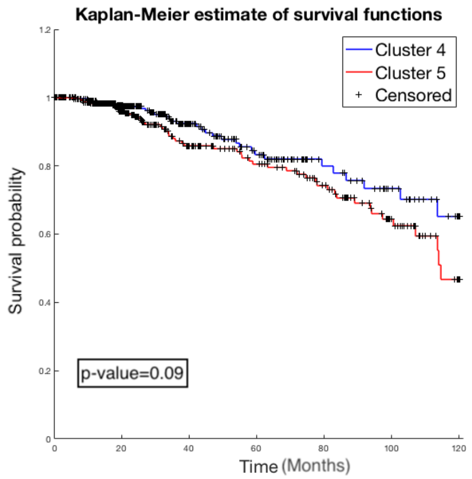


(b)

**Figure 7.** (a) Hierarchical clustering of 726 samples using only gene expression of 290 genes. Even though the difference in survival counts between two major clusters (4 and 5) is still significant (chi-squared test's p-value= 0.02), it is not as strongly separated as in the integrative method. (b) Graphical display of survival rate using the Kaplan-Meier curves. Sample survival times (months) are plotted on the x-axis (truncated at 10 years), and the probability of survival calculated according to the Kaplan-Meier method is plotted on the y-axis. After excluding copy number alteration and methylation, the Kaplan-Meier curves of major clusters do not have a significant survival rate difference where the log-rank's p-value is 0.09.

**Additional data table S1.** The list of 150 significant genes chosen after performing iWCluster of TCGA breast cancer data. We used ANOVA to choose 150 top genes (from 3,426 genes in the study) that have significantly different mean values of integrative measures among the four clusters. We then sorted these 150 genes based on the highest to lowest mean values in cluster 1.

| | | | | |
|---|---|---|---|---|
| 1. ERBB2 | 31. JAM2 | 61. ST5 | 91. GNA11 | 121. LEPR |
| 2. GRB7 | 32. PRDM2 | 62. DHH | 92. AMH | 122. FEZ1 |
| 3. PIK3C2A | 33. TLR4 | 63. IGFBP6 | 93. KLF4 | 123. MED31 |
| 4. RND1 | 34. IDE | 64. ZBTB17 | 94. HDAC7 | 124. FSTL3 |
| 5. SH3BGRL3 | 35. STX12 | 65. SIPA1L1 | 95. EHD2 | 125. TP73 |
| 6. ERBB3 | 36. RB1 | 66. PTCH2 | 96. SYTL1 | 126. AOC3 |
| 7. HSP90B1 | 37. PLA2G2A | 67. ATM | 97. TP53BP1 | 127. IL4R |
| 8. TCAP | 38. KLF9 | 68. BTRC | 98. ZBTB16 | 128. VAMP2 |
| 9. IKZF3 | 39. ACVRL1 | 69. JDP2 | 99. SORBS1 | 129. EEF2 |
| 10. ITGA5 | 40. TGFB3 | 70. RAPGEF3 | 100. CIDEA | 130. FZD2 |
| 11. MED24 | 41. COQ6 | 71. JAM3 | 101. NFE2 | 131. SERPINF2 |
| 12. THRA | 42. CRB2 | 72. DLG4 | 102. HHEX | 132. PRTN3 |
| 13. MED1 | 43. NAA16 | 73. KIF17 | 103. RPL26 | 133. NLRP1 |
| 14. ITGB4 | 44. ACTA2 | 74. HSPB2 | 104. PER3 | 134. XPA |
| 15. RORA | 45. PNRC2 | 75. NDOR1 | 105. UBR1 | 135. STAT5B |
| 16. CDK12 | 46. POLR2A | 76. HOXC8 | 106. PIP5K1C | 136. SSTR3 |
| 17. CASC3 | 47. EVL | 77. RAB27A | 107. GRIN3B | 137. EGR2 |
| 18. PIP4K2B | 48. TCF12 | 78. ILK | 108. ATP1B2 | 138. TNFSF13 |
| 19. SAP30BP | 49. BIRC3 | 79. CUEDC2 | 109. NDN | 139. TRIM22 |
| 20. ABL1 | 50. SVEP1 | 80. NFIC | 110. PER1 | 140. UBXN11 |
| 21. PTEN | 51. TNFSF12 | 81. STAT6 | 111. DOCK8 | 141. PIK3CD |
| 22. SMARCD1 | 52. CASP7 | 82. APBB1 | 112. CRY2 | 142. PRKCB |
| 23. BNC2 | 53. REEP6 | 83. FAM160A2 | 113. KLK4 | 143. ETS1 |
| 24. AMHR2 | 54. MVP | 84. RHOJ | 114. STX8 | 144. RHOG |
| 25. ZFP36L1 | 55. MAP1A | 85. THBS1 | 115. DLEU1 | 145. RAB11B |
| 26. CRK | 56. RPS6KA5 | 86. TJP3 | 116. GJC1 | 146. JUN |
| 27. MRVI1 | 57. ITM2B | 87. CASP9 | 117. CBL | 147. STAT3 |
| 28. GRK5 | 58. VPS11 | 88. PGR | 118. EPS15 | 148. GNG7 |
| 29. GUCY1A2 | 59. FAR1 | 89. CLIC6 | 119. SLC2A4 | 149. CTSG |
| 30. NEDD4 | 60. PAFAH1B2 | 90. HSPG2 | 120. TGFBR3 | 150. STAT5A |

**Additional data table S2.** The list of 166 genes associated to the cluster with the lowest survival rate. These genes have significantly different mean values in this cluster compared to other three clusters using Bonferroni corrected p-value of 0.01 after t-test.

| | | | |
|---|---|---|---|
| 1. ACTN4 | 43. FOXO1 | 85. PIGU | 127. SCYL1 |
| 2. ACVRL1 | 44. FSTL3 | 86. PIP5K1C | 128. SERPINF2 |
| 3. AKR1A1 | 45. FZD2 | 87. PITPNM2 | 129. SHARPIN |
| 4. AMH | 46. GABPA | 88. PLK3 | 130. SHKBP1 |
| 5. AMHR2 | 47. GGA3 | 89. PMCH | 131. SIGLEC11 |
| 6. ANGPTL3 | 48. GJC1 | 90. PMM1 | 132. SLC22A11 |
| 7. APIP | 49. GNA11 | 91. PNRC2 | 133. SLC4A8 |
| 8. ARL2 | 50. GPAA1 | 92. POP1 | 134. SLC9A5 |
| 9. ARRB1 | 51. GPBP1L1 | 93. PPFIA1 | 135. SPEN |
| 10. ASAP1 | 52. GTPBP3 | 94. PPP1R16A | 136. SS18L1 |
| 11. ASF1B | 53. HDHD2 | 95. PPP2R2D | 137. STAT3 |
| 12. ATF1 | 54. HHEX | 96. PRDM2 | 138. STAT6 |
| 13. ATM | 55. HSF1 | 97. PRKCB | 139. STX12 |
| 14. BACH1 | 56. HSPB7 | 98. PRKCH | 140. SUMO2 |
| 15. BIRC3 | 57. IDE | 99. PRMT5 | 141. SUPT5H |
| 16. BPTF | 58. IGF1 | 100. PRPF6 | 142. SYTL1 |
| 17. CABLES2 | 59. IL4R | 101. PRPSAP1 | 143. TALDO1 |
| 18. CASP9 | 60. ITGB4 | 102. PSMA7 | 144. TCF12 |
| 19. CDCA5 | 61. ITM2B | 103. PSMB3 | 145. TCF3 |
| 20. CDH24 | 62. ITPK1 | 104. PSMB5 | 146. TGFB3 |
| 21. CEP250 | 63. JUN | 105. PSMC1 | 147. TGFBR3 |
| 22. CHGB | 64. KCNG1 | 106. PSMC4 | 148. TGIF1 |
| 23. CHRM4 | 65. KEAP1 | 107. PSMC5 | 149. THBS1 |
| 24. CLIC6 | 66. KLF4 | 108. PSMD12 | 150. TIMM50 |
| 25. CLTA | 67. LCP1 | 109. PSMD13 | 151. TJP3 |
| 26. CLTC | 68. LOH12CR1 | 110. PSMD8 | 152. TOM1 |
| 27. CPNE1 | 69. MAF1 | 111. PSTPIP1 | 153. TOM1L1 |
| 28. CREBBP | 70. MCOLN1 | 112. PUF60 | 154. TRIM22 |
| 29. DCAF7 | 71. MED17 | 113. PWP2 | 155. TRPC4AP |
| 30. DLG4 | 72. MED30 | 114. RAB6A | 156. TXNL4B |
| 31. EFS | 73. MFSD3 | 115. RABGAP1 | 157. UBE2M |
| 32. EIF3G | 74. MOCS3 | 116. RAPGEF3 | 158. UBE2O |
| 33. EIF3H | 75. MRI1 | 117. RB1 | 159. UBR1 |
| 34. EIF3K | 76. MYT1 | 118. RECQL4 | 160. USP6NL |
| 35. EIF6 | 77. NEDD4 | 119. REEP6 | 161. VPS33B |
| 36. EMP3 | 78. NFE2 | 120. RHOG | 162. WDR24 |
| 37. EPS15 | 79. NLRP1 | 121. RPL30 | 163. WDYHV1 |
| 38. ETS1 | 80. NOS2 | 122. RPL8 | 164. ZBTB16 |
| 39. EXOSC1 | 81. P4HA3 | 123. RPP30 | 165. ZBTB17 |
| 40. EXOSC4 | 82. PAAF1 | 124. RPS3 | 166. ZBTB5 |
| 41. FAM188A | 83. PAX1 | 125. RYR1 | |
| 42. FAM60A | 84. PDHX | 126. SAP30BP | |
